# *Escherichia coli* aggregates mediated by native or synthetic adhesins exhibit both core and adhesin-specific transcriptional responses

**DOI:** 10.1101/2023.02.14.528454

**Authors:** Yankel Chekli, Rebecca J. Stevick, Etienne Kornobis, Valérie Briolat, Jean-Marc Ghigo, Christophe Beloin

## Abstract

Bacteria can rapidly tune their physiology and metabolism to adapt to environmental fluctuations. In particular, they can adapt their lifestyle to the close proximity of other bacteria or presence of different surfaces. However, whether these interactions trigger transcriptomic responses is poorly understood. We used a specific set up of *E. coli* strains expressing native or synthetic adhesins mediating bacterial aggregation to study the transcriptomic changes of aggregated compared to non-aggregated bacteria. Our results show that following aggregation, bacteria exhibit a core response independent of the adhesin type, with differential expression of 56.9% of the coding genome, including genes involved in stress response and anaerobic lifestyle. Moreover, when aggregates were formed via a naturally expressed *E. coli* adhesin (Antigen 43), the transcriptomic response of the bacteria was more exaggerated compared to aggregates formed via a synthetic adhesin. This suggests that the response to aggregation induced by native *E. coli* adhesins could have been finely tuned during bacterial evolution. Our study therefore provides insights on the effect of self-interaction in bacteria and allows a better understanding of why bacterial aggregates exhibit increased stress tolerance.

**Importance:** Formation of bacterial aggregates has an important role in both clinical and ecological contexts. Although these structures have been previously shown to be more resistant to stressful conditions, the genetic basis of this stress tolerance associated with the aggregate lifestyle is poorly understood. Surface sensing mediated by different adhesins can result in varying changes on bacterial physiology. However, whether adhesin-adhesin interactions as well as the type of adhesin mediating aggregation affects bacterial cell physiology is unknown. By sequencing the transcriptomes of aggregated and non-aggregated cells expressing native or synthetic adhesins, we characterized the effects of aggregation and adhesin type on *E. coli* physiology.

## Introduction

Bacteria often live in communities called biofilms, adhering to each other and to biotic or abiotic surfaces. Auto-aggregation, defined as bacterium-bacterium interactions of genetically identical strains (Trunk et al., 2018), is observed in environmental and pathogenic species, and contributes to biofilm formation (Kragh et al., 2016; Sorroche et al., 2012). The importance of aggregate formation has increased due to the awareness that, in most cases, natural and clinical biofilms resemble cell aggregates rather than large highly structured biofilms (Bjarnsholt et al., 2013; Sauer et al., 2022). Although rarely larger than a few hundred micrometers, these aggregates maintain biofilm characteristics, including enhanced tolerance to physical and chemical stresses and host immune defenses (Alhede et al., 2020; Blom et al., 2010; Haaber et al., 2012). In *Pseudomonas aeruginosa* aggregation is induced upon exposure to detergent (Klebensberger et al., 2006) and was shown to promote *P. aeruginosa* survival to antimicrobial treatments in patients with cystic-fibrosis (Caceres et al., 2014; Walker et al., 2005). Moreover, while *P. aeruginosa*, and other species of bacteria such as *Sphingobium*, use aggregation as a defense mechanism against predation by protozoa (Blom et al., 2010; Matz et al., 2004), the formation of aggregates was also shown to be an essential step that promotes *Neisseria meningitidis* virulence (Bonazzi et al., 2018).

*Escherichia coli* a versatile bacterium that can behave either as a pathogen or a commensal, is prone to aggregation mediated by a range of adhesins, including the autotransporter Antigen 43 (Ag43). The self-recognition properties of Ag43 promote auto-aggregation and biofilm formation (Ageorges et al., 2019; Henderson et al., 1997; Ulett et al., 2007). Ag43-dependent aggregates exhibit specific properties including protection from neutrophil killing, promotion of intracellular aggregate and biofilm formation, contributing to persistence and virulence of uropathogenic *E. coli* in the mouse bladder (Ulett et al., 2007).

While it is well-accepted that aggregate formation correlates with stress tolerance, the underlying mechanisms allowing aggregated bacteria to grow or survive under adverse conditions remain unknown. Here, we used RNA-seq and a specific setup of *E. coli* K12 strains expressing either native (Ag43) or nanobody-based synthetic self-recognizing adhesins to identify changes in gene expression associated with stress resistance between non-aggregated and aggregated cells. We found that, regardless of the adhesin mediating aggregation, the aggregates exhibit a core transcriptomic signature indicative of a specific metabolic rewiring and characterized by the activation of several stress resistance pathways. Beyond this core response to aggregation, we also showed that many genes were specifically regulated in Ag43-dependent aggregates compared to aggregates mediated by synthetic adhesins. These results demonstrate the existence of a specific physiological response to aggregation and suggest that aggregate physiology leads to profound metabolic changes as well as the activation of multiple stress response systems.

## Results

### Both native and synthetic adhesins promote equivalent *E. coli* auto-aggregation and tolerance to antibiotic stress

To better understand the impact of auto-aggregation on *E. coli* physiology, we chose a combination of native and “synthetic” self-recognizing adhesins to investigate both general and adhesin-specific transcriptomic responses. Ag43 (also known as Flu) was used as a native adhesin since it is the major self-recognition and aggregation factor in *E. coli* K12. Ag43 is a naturally phase variable adhesin being either in an ON (Ag43+) or OFF (no Ag43) status. Ag43 ON and OFF status was monitored through the use of an operon fusion between *ag43* and the *lacZ* gene (Chauhan et al., 2013). For this adhesin, we used a series of four strains (**Fig. 1A**). Two strains represent the aggregated state: i) PcL*ag43*, constitutively expressing Ag43 and ii) Ag43_On containing cells in an Ag43 ON state with native levels of Ag43. Two other strains represent the non-aggregated state: i) Δ*ag43* deleted for *ag43* (no Ag43) and ii) Ag43_Off containing cells in an Ag43 OFF state.

**Figure 1.**
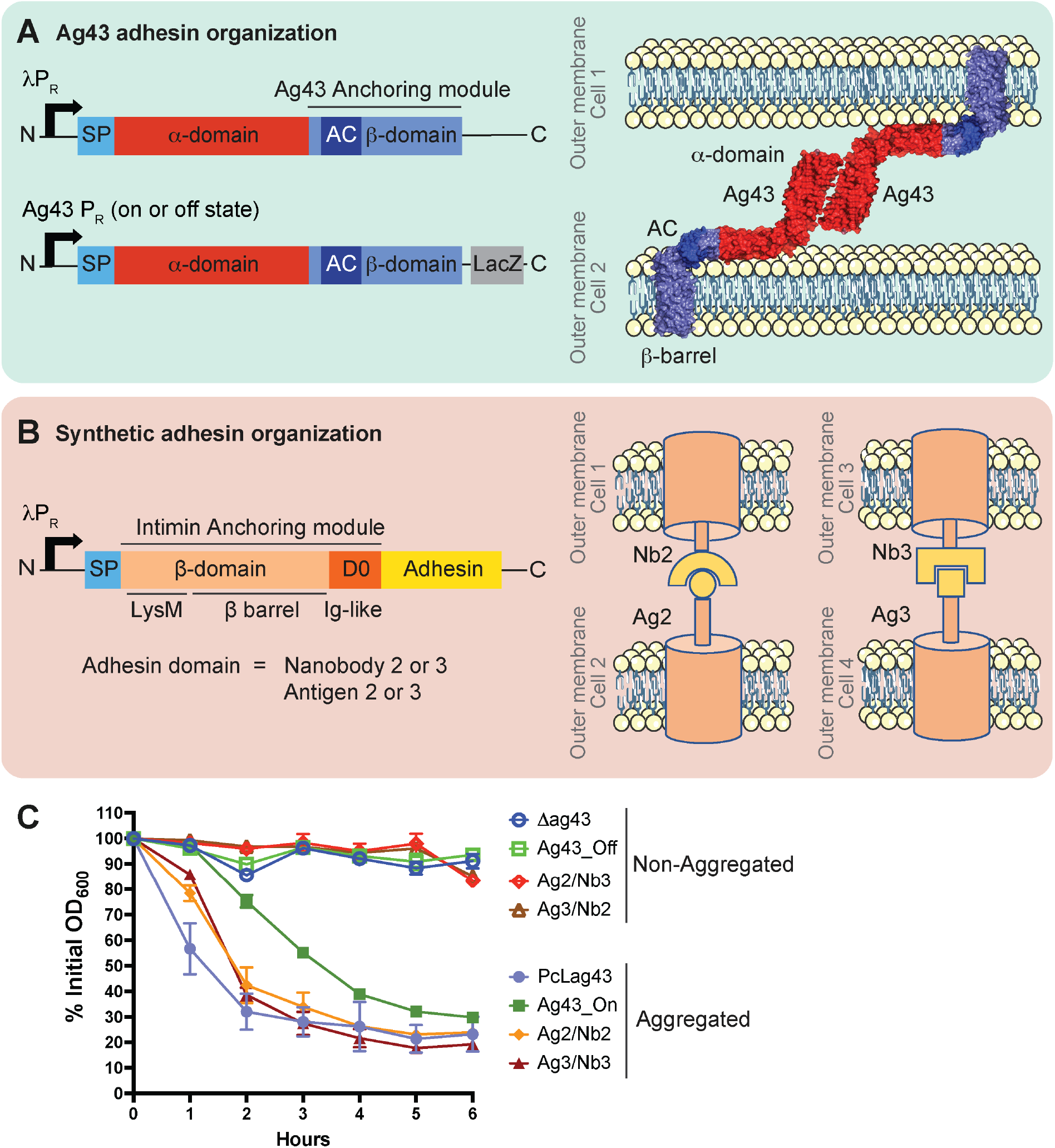
Genetic organization and aggregation capacity of the adhesins used in this study. **(A)** Genetic organization of the native Ag43 adhesin constructions, SP= signal peptide, AC = Autochaperone domain. **(B)** Genetic organization of the synthetic adhesin constructions, SP= signal peptide. In the different synthetic adhesin strains, the yellow part corresponds to Nanobody 2, or its corresponding Antigen 2, or Nanobody 3 or its corresponding Antigen 3.**(C)** Aggregation curves realized with all the strains and strain couples used in this study. Results are expressed as a percentage of the OD_600_ measured at the top of the tube at T0. For each strain a measurement is taken at the exact same position in the tube throughout time. A higher % OD measurement corresponds with no aggregation and a decrease in % OD indicates aggregate formation. Plotted data represents the mean ± standard deviation of 3 biological replicates (each replicate is the mean of 2 technical replicates).

For the synthetic adhesins, we used chimeric fusion proteins consisting of an anchoring and an exposed domain (Glass & Riedel-Kruse, 2018) (**Fig. 1B**). The outer membrane anchor is composed of the intimin N terminus from an enterohemorrhagic *E. coli* (EHEC O157:H7), including a small N-terminal signal peptide, a LysM domain for peptidoglycan binding, and a β -barrel for transmembrane insertion (Glass & Riedel-Kruse, 2018; Salema et al., 2013). The exposed domains, composed of a nanobody (the variable domains of camelid heavy-chain antibodies) or its corresponding antigen, were then fused to the C-terminus part of the truncated intimin. Recognition between a nanobody (Nb2 or Nb3) expressed on one *E. coli* cell and its corresponding antigen (Ag2 or Ag3) on another cell promotes aggregation. We swapped the *ag43* gene for the different synthetic adhesins genetic constructions and placed them under the control of the constitutive lambda P_R_ promoter PcL. For the synthetic adhesins, the aggregated state is represented by a 1:1 mixture of cells expressing Nb2 and Ag2, or Nb3 and Ag3. The non-aggregated state is represented by a 1:1 mixture of cells expressing Nb2 and Ag3, or Nb3 and Ag2 **(Fig. 1B)**.

We first verified the capacity of each adhesin to promote aggregation. After 6 hours of incubation, strains PcL*ag43* and Ag43_On, and cultures composed of Ag2/Nb2 and Ag3/Nb3 provided similar levels of aggregation, while the Δ*ag43*, Ag43_Off strains, and cultures composed of Ag2/Nb3 and Ag3/Nb2 displayed no aggregation **(Fig. 1C)**. Measurements of the aggregate sizes in culture also revealed that although the size of Ag2/Nb2, Ag3/Nb3 and Ag43_On aggregates were comparable, aggregates made with PcL*ag43* strains were almost 10 times bigger, suggesting that overexpressing Ag43 led to stronger interactions resulting in an increased aggregate size. **(Supplementary Fig. S1)**. We then assessed the capacity of the different aggregated cells to sustain a lethal antibiotic stress as compared to their non-aggregating counterparts. Aggregated cells survived the lethal action of amikacin much better than the non-aggregated cells, therefore validating our set-up to evaluate the corresponding aggregate physiology (**Supplementary Fig. S2)**.

### Auto-aggregation leads to a robust core transcriptional response independently of the adhesin type

In order to perform comparative RNAseq, we developed a protocol to optimize the separation between planktonic cells and aggregates. For this, different strains and synthetic mixes were independently cultured in LB until OD = 0.5 and then left to aggregate in separating funnels for 3 hours at 37°C in static conditions (**Supplementary Fig. S3**). One mL of cells was sampled from the bottom of each funnel to perform RNA extraction and a comparative transcriptomic analysis of the different types of cultures.

The overall transcriptional profiles of each condition clustered depending on the strains’ capacity to aggregate and the type of adhesin expressed (Ag43 natural or nanobodies synthetic), as shown in the PCA calculated using normalized transcript counts (**Fig. 2**). The first principal component (PC1), explaining 38% of the variance, separates the samples based on aggregation, regardless of the type of adhesin. Native and synthetic aggregates clustered along the x-axis, with non-aggregated controls also clustered based on PC1. On the other hand, the second principal component (PC2), explaining 33% of the variance, separates the samples based on their adhesin type, with the samples expressing native adhesins being very close on the y-axis and separate from the bacteria expressing synthetic adhesins. Thus, while we can see that the aggregates, native and synthetic, are close to each other according to PC1, suggesting a core response of the bacteria towards aggregation, we can also observe that the samples are different according to PC2. This illustrates that, in addition to aggregation, the type of adhesin used, native or synthetic, is a discriminating factor, suggesting a specific response of bacteria depending on the adhesin.

**Figure 2.**
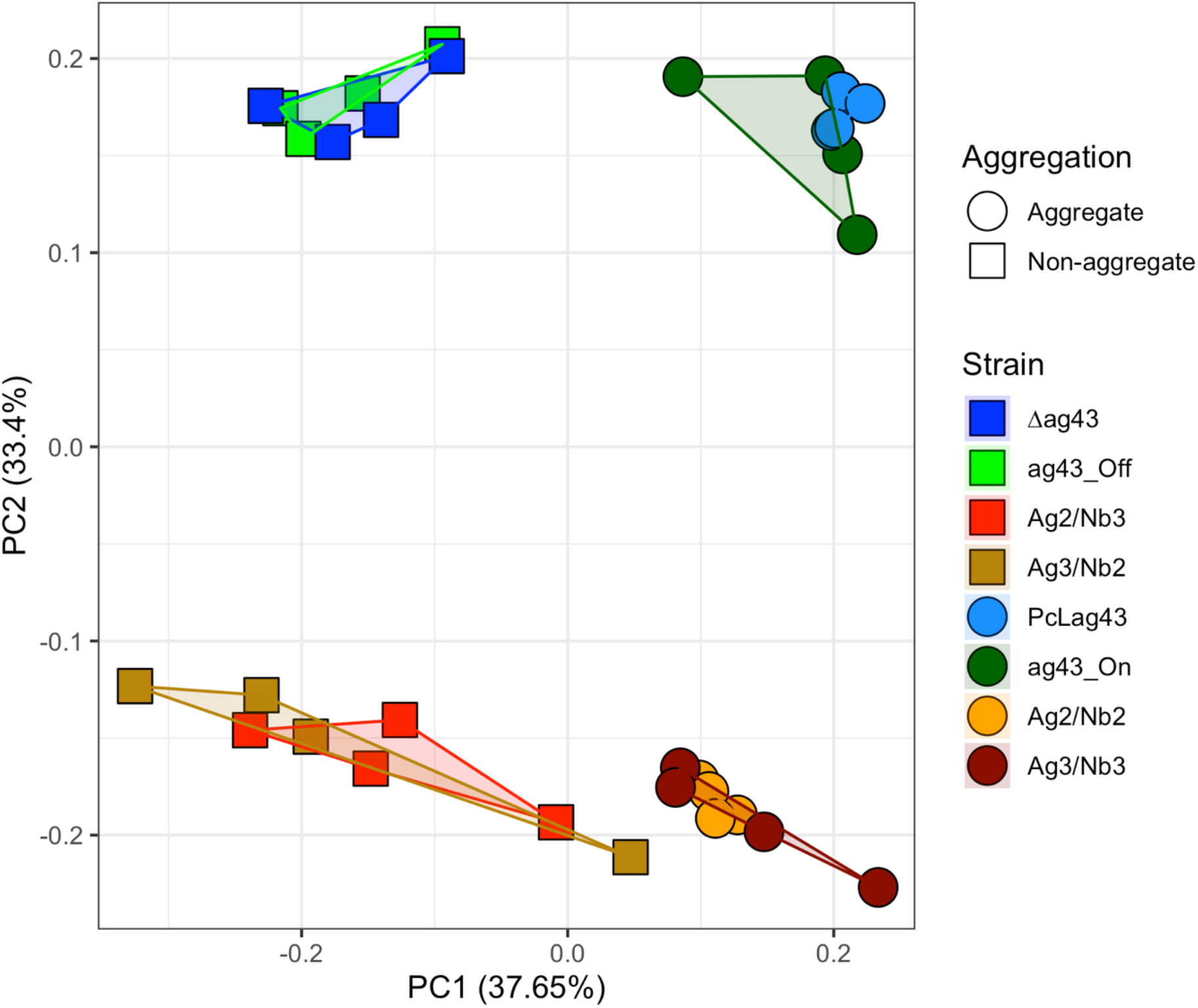
Aggregation capacity and the adhesin type affect overall transcriptional profiles. Principal component analysis (PCA) of all samples sequenced (4 biological replicates per each of the 8 conditions) based on Variance Stabilizing Transformation (VST) normalized transcript counts. Colors represent the different strains while shapes represent the aggregation status.

The *E. coli* K-12 genome has 4,401 genes encoding 116 RNAs and 4,285 proteins (Serres et al., 2001). In order to compare between non-aggregates and aggregates, differential expression analysis was conducted for each non-aggregating strain to the 2 aggregating strains of the same adhesin type (Native or synthetic) (**Fig. 3, barplot on the left side**). Considering these 8 different comparisons independently, we found that the “pan” response during aggregation is composed of 2,504 genes (56.9% of all coding genes) that were significantly differentially expressed upon aggregation for any adhesin type (considered as significant when both Log2-fold change ≥ 1 or ≤ − 1 and adjusted p-value < 0.05) (**Supplementary Table S1**). This indicates that aggregation leads to a profound transcriptional reprogramming of the bacteria. Of these 2,504 genes, 1380 were downregulated (31.3% of all coding genes) and 1147 were upregulated (26.0%) (23 genes were either up or down regulated depending on the comparison).

**Figure 3.**
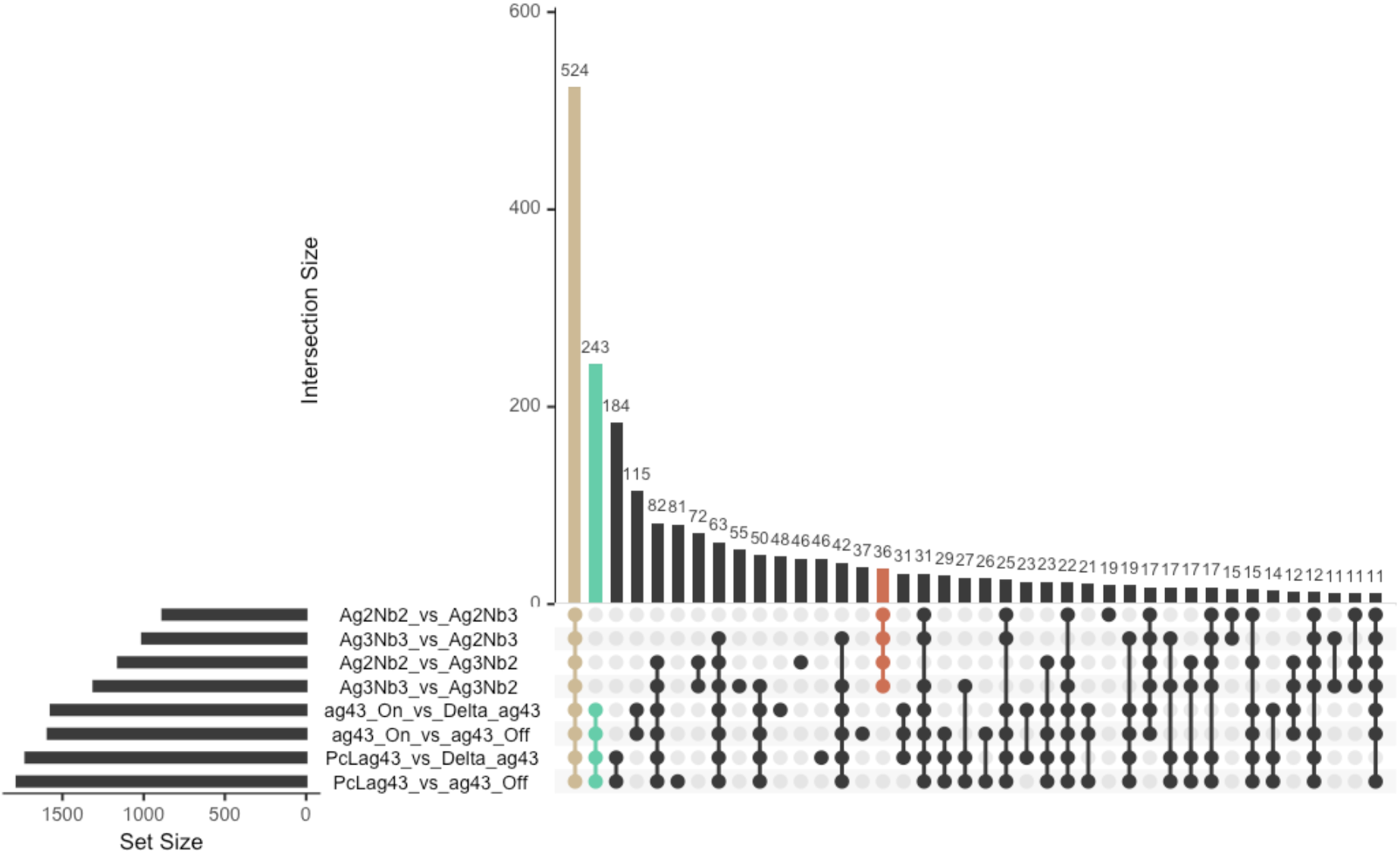
The core transcriptomic response of bacteria to aggregation is independent of the type of adhesin mediating aggregation. Upset plot (Venn diagram alternative) showing the number of significantly differentially expressed genes shared between each comparison (vertical bars). The total number of genes significantly differentially expressed in each comparison is shown in the horizonal bar graph on the left. In beige the genes differentially regulated as the core aggregate vs non-aggregate response. In green the genes differentially regulated specifically in Ag43-mediated aggregates. In red the genes differentially regulated specifically in Nanobody-mediated aggregates. Genes were considered significant when Log2-fold change ≥ 1 or ≤ − 1 and adjusted p-value < 0.05.

From these 8 different comparisons, we then identified the “core” response of *E. coli* towards aggregation, *i*.*e*. the genes that were up or down regulated in all of the 8 comparisons of aggregates vs non-aggregates. This aggregation “core” response contained 524 genes (11.9% of the coding genes) (208 upregulated, 4.7% and 316 downregulated, 7.2%) (**Fig. 3, beige bar** and **Supplementary Table S2**) and corresponded to the group sharing the highest number of significantly regulated genes. This emphasized that there is a common response of bacterial aggregates formed upon expression of native and synthetic aggregates.

### Aggregation leads to profound changes in metabolism and a global downshift of essential cellular functions

To characterize the global effects of aggregation on biological processes, we used the PANTHER bioinformatics web server (http://www.pantherdb.org) (The Gene Ontology, 2019) to determine the Gene Ontology (GO) groupings that were over or under-represented among significantly regulated genes in the core aggregate vs non-aggregate response. According to the number of significant genes, the over-representation assay determined if the pathway is significantly different than would be expected by chance. The result is expressed as fold enrichment. Multiple pathways were found to be significantly over-represented (26 pathways) or under-represented (36 pathways) in aggregates compared to non-aggregates (**Fig. 4 and Supplementary Table S3**, see also Supplementary Text for more detailed information).

**Figure 4.**
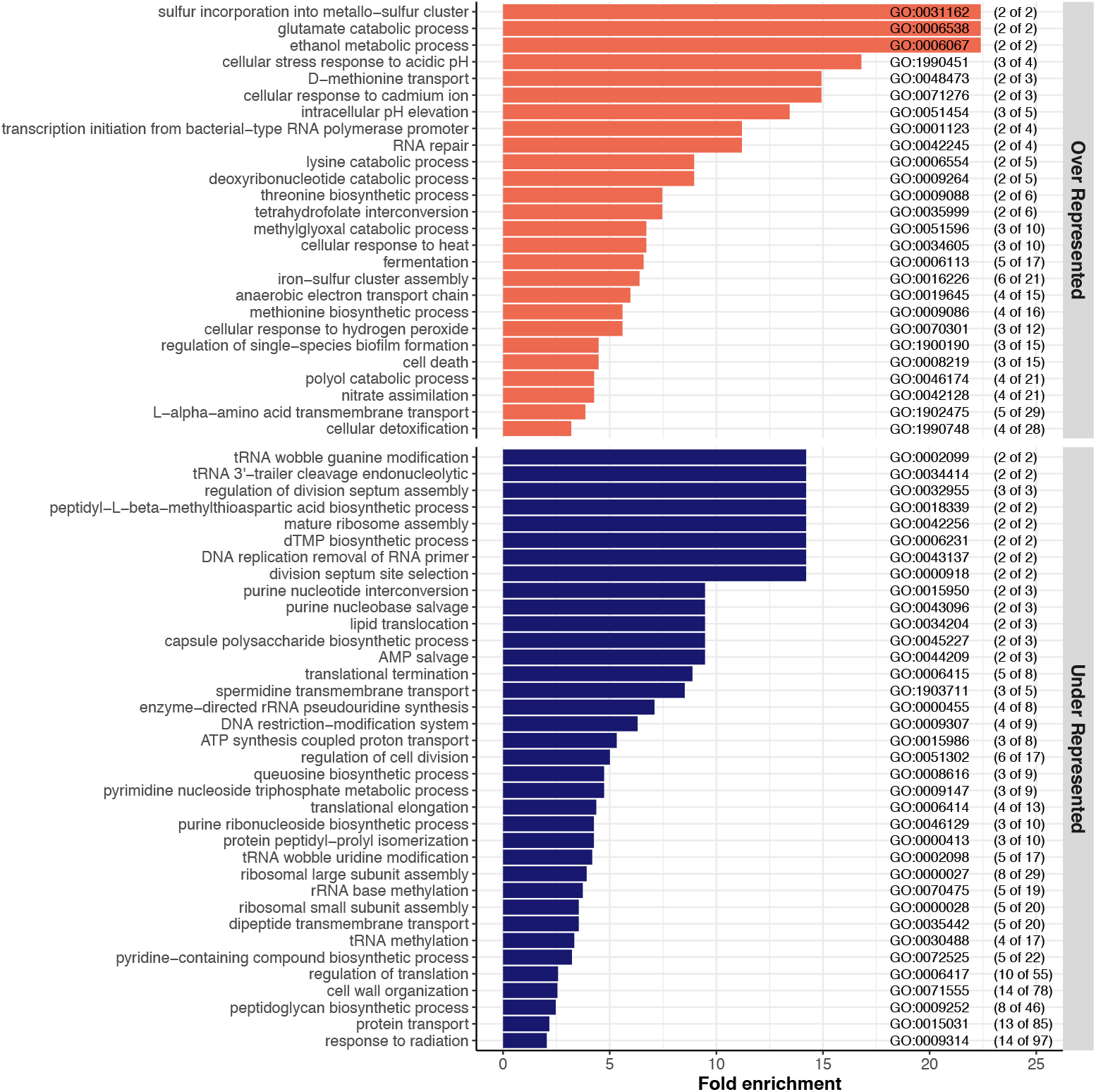
Visualization of the GO biological processes significantly over- and under-represented in all aggregate vs non-aggregate comparisons. The upper panel represents the over-represented pathways (orange bars) and the bottom panel represents the under-represented pathways (blue bars). The GO enrichment analysis has been performed via PANTHER using the biological process (BP) category. The genes included in this analysis are the 524 genes commonly regulated in all the comparisons of aggregates vs non-aggregates (beige bar in Fig 3). The GO term for each biological process pathway is indicated on the right. The numbers in parentheses indicate the number of genes significantly up/downregulated in the aggregates compared to non-aggregated cells out of the total number of genes in the pathway.

### Anaerobic metabolism

Several of the over-represented pathways were linked to a switch from aerobic to anaerobic metabolism (pink highlighted pathways in **Supplementary Table S3**). For example, we found that upon aggregation, pathways related to fermentation, anaerobic respiration chains and nitrite reduction were significantly enriched. Some genes classically induced under anaerobia (or microaerophilic conditions) and indicative of amino-acid metabolism modification were also induced upon aggregation, such as *adhE* that catalyzes the reduction of acetyl-coA to ethanol, and *cadAB* and *lcdC*, which are involved in lysine decarboxylase pathways.

#### Amino acid metabolism, nucleotide production and cell growth

We detected additional modification of amino-acid metabolism with the most over-represented pathways corresponding to methionine biosynthesis/transport, threonine biosynthetic process, and glutamate catabolic process (yellow highlighted pathways in **Supplementary Table S3**). This reduction of essential growth functions is further supported by the downregulation of functions linked to ribosome assembly and functioning (green highlighted pathways in **Supplementary Table S3**), DNA replication and division (orange highlighted pathways in **Supplementary Table S3**) and pathways involved in the synthesis and/or salvation of purines as well as pyrimidine synthesis. This decrease in nucleotide synthesis genes and recycling of nucleotides as well as the downregulation of both RNase HI and HII could lead to a decrease in the fidelity and the rate of DNA replication and therefore a decrease in the growth rate of bacteria. Additionally, three pathways involved in the regulation of cell division, positioning and assembly of the division septum were under-represented. Finally, small and large ribosomal subunit assembly pathways were under-represented in aggregates along with ribosome assembly factors and RNA helicases. This reduction of ribosomal function might also be linked to induction of stringent response with multiple (p)ppGpp alarmone activated genes.

#### Modification of cell envelope

Cell envelope synthesis and especially peptidoglycan (PG) synthesis pathways were greatly impacted by aggregation (violet pathways in **Supplementary Table S3**). This is consistent with an under-representation of pathways involved in membrane expansion that also indicate decreased metabolism, growth and division. For instance, several pathways related to the cell wall were under-represented: lipid translocation, capsule polysaccharide biosynthetic process, cell wall organization, peptidoglycan biosynthetic process and protein transport. This supports the important modification of PG biogenesis and turnover that might be present in aggregated cells. In addition to PG related genes, four out of the seven genes involved in the lipoprotein posttranslational modification pathway were downregulated (*lgt, ispA, lolC* and *lolD*) (see **Supplementary Table S2**) also suggesting a modification of the lipoprotein composition of the membranes upon aggregation.

#### Biofilm-related genes

Since aggregates can be considered as early-stage biofilms, we also focused on genes known to be involved in biofilm phenotypes. Some of these were among the most upregulated genes in aggregates: *bhsA, bssS, bssR*, and *bolA. bhsA* was shown to be induced by multiple stresses and it seems to promote or reduce biofilm formation depending on the growth medium (Beloin et al., 2004; Zhang et al., 2007). Expression of *bssS* and *bssR* was also shown to be increased in biofilms (Ren et al., 2004) and their associated proteins repress motility of cells in LB, the medium used in our study (Domka et al., 2006). Therefore, increased expression of these genes may lead to reduced motility upon aggregate formation to avoid dispersal. Consistently, *bolA*, a motile/adhesive transcriptional switch involved in the transition from the planktonic state to a biofilm state, is also induced in aggregates (Dressaire et al., 2015).

Taken together, these results indicate that aggregation leads to a profound metabolic rewiring together with reduction of several essential functions, indicating a global reduction of growth.

### Aggregates transcriptionally respond to different stresses

The reduction of growth upon aggregate formation could be caused by stressful conditions developing within the aggregates. Accordingly, many genes related to various stress responses were induced upon aggregation (pathways highlighted in red in **Supplementary Table S3**). Two out of four acid resistance systems, GadABC and CadAB/LcdC, along with other acid stress response genes (*yhiM, hdeAB, hchA, dps*) were enriched in aggregates. We also identified several genes associated with oxidative stress like hydrogen peroxide (*ychH, ygiW* and *katG)* and cellular detoxification (*katG, sodB, hcp* and *frmA)* or protection (*dps*) that were significantly over-expressed in aggregates. This strengthens the hypothesis that certain bacteria within the aggregates maintain an aerobic metabolism. In addition to acidic or oxidative stress responses, we found several up-regulated genes involved in response to heat (*clpA, clpB, rpoH*) and general stress response (including *rpoS* and *rpoS*-activated genes). Finally, multiple genes that are induced or repressed by (p)ppGpp were identified as significantly regulated upon aggregation, suggesting an increased (p)ppGpp level from starvation and activation of the stringent response. These results indicate that cells forming aggregates were subjected to various types of stresses and activate multiple pathways to counteract them.

### Aggregation response is associated with antibiotic tolerance

Consistent with previous studies, we determined that aggregated cells displayed enhanced survival when exposed to a lethal antibiotic stress (**Supplementary Fig. S2)**. While this increased tolerance could be linked to reduced growth and the different stress responses induced, we detected at least three other specific systems that were impacted in aggregates compared to non-aggregated cells and could be involved in sustaining antibiotic stress: tRNAs modification and processing, toxin-antitoxin systems and iron-sulfur systems.

#### tRNAs modification and processing

Four different pathways linked with modification and processing of tRNAs were under-represented in aggregates: tRNA wobble guanine modification, tRNA wobble uridine modification, tRNA 3’-trailer cleavage, and tRNA methylation (blue highlighted in pathways in **Supplementary Table S3**). Additionally, other tRNA modification genes as well as numerous genes encoding proteins that modify either 16S RNA, 23S RNA or ribosomal proteins were also downregulated in aggregates (see **Supplementary Table S2**). Decreased expression of these different translational modifiers might be increasing the rate of mistranslation and thus enhanced tolerance towards several antibiotics (Witzky et al., 2019). Notably some recent works have shown that specific tRNA modification could be directly involved in enhanced tolerance to aminoglycosides (Babosan et al., 2022; Fruchard et al., 2022).

#### Toxin-antitoxin systems

We found that pathways involved in cell death and production of toxic compounds were enriched upon aggregation, as well as upregulation of genes encoding toxins belonging to toxin-antitoxin systems (TA). Since some of the toxins of these TA systems can act on different essential cellular processes such as cell wall synthesis, membrane integrity, replication, transcription, translation or cytoskeleton formation, they might modify the tolerance of bacteria towards antibiotics, although formal demonstration of such effects are still missing (Dörr et al., 2010; Nathan Fraikin et al., 2019; N. Fraikin et al., 2019; Jurėnas et al., 2022; Kim et al., 2009; Korch & Hill, 2006; Singh et al., 2021; Soo & Wood, 2013; Unterholzner et al., 2013; Vazquez-Laslop et al., 2006; Yang & Walsh, 2017).

#### Iron-sulfur cluster systems

We observed that the Suf system is induced upon aggregation (*sufABCDSE*). *E. coli* uses two systems to assemble iron-sulfur clusters: the Ics and the Suf systems (Garcia et al., 2019). The Ics system is used under normal growth conditions while the Suf system is employed in condition of iron-depletion or oxidative stress. The *suf* genes were induced by superoxide generators and hydrogen peroxide (Lee et al., 2004), further indicating that aggregated cells are subjected to multiple stresses. Interestingly, cells using the Suf system display enhanced tolerance towards aminoglycosides through a reduced proton motive force (pmf), decreasing entry of the antibiotics into the cells (Ezraty et al., 2013). This mechanism might be at play to explain the enhanced amikacin tolerance observed in our aggregates (**Supplementary Fig. S2**).

Altogether, these results demonstrate the existence of a robust and mainly anoxia- and stress-related transcriptional response upon aggregation independently of the type of adhesin.

#### Aggregation mediated by native *E. coli* adhesin leads to a specific transcriptional response with enhanced anaerobia and cell growth signatures

While we identified a robust core transcriptomic response upon aggregation, we also detected responses specific to the adhesin used to mediate aggregation (**see Figure 2**) by comparing all the genes that were differentially regulated only in the native Ag43-mediated aggregates or only in the synthetic nanobody-mediated aggregates (**Fig. 3** green and red bars). Clustering the genes using the GO database and enrichment analysis allowed us to identify differences in regulatory pathways between the two modes of aggregation.

In aggregates mediated by *E. coli* native Ag43 adhesin, we found that 243 genes (5.6% of coding genes) were uniquely up or down regulated when compared to non-aggregated controls (**Fig. 3**, red bar). Out of these 243 genes, 122 were upregulated (2.8%) and 121 were downregulated (2.8%) (**Fig. 5 representing the pathways associated with these genes** and **supplementary Table S4** and **S5**). For the nanobody-mediated aggregates, the number of specific and differentially regulated genes was much smaller, with only 36 genes (0.8%) that were up-or downregulated compared to non-aggregated control (**Fig. 3**, green bar). Of these 36 genes, 17 were upregulated (0.38%) and 19 were downregulated (0.43%) (**Fig. 6 representing the pathways associated with these genes** and **supplementary Table S6** and **S7**). By contrast, the specific transcriptomic signature of nanobodies-mediated aggregation was limited, especially compared to the core response to aggregation.

**Figure 5.**
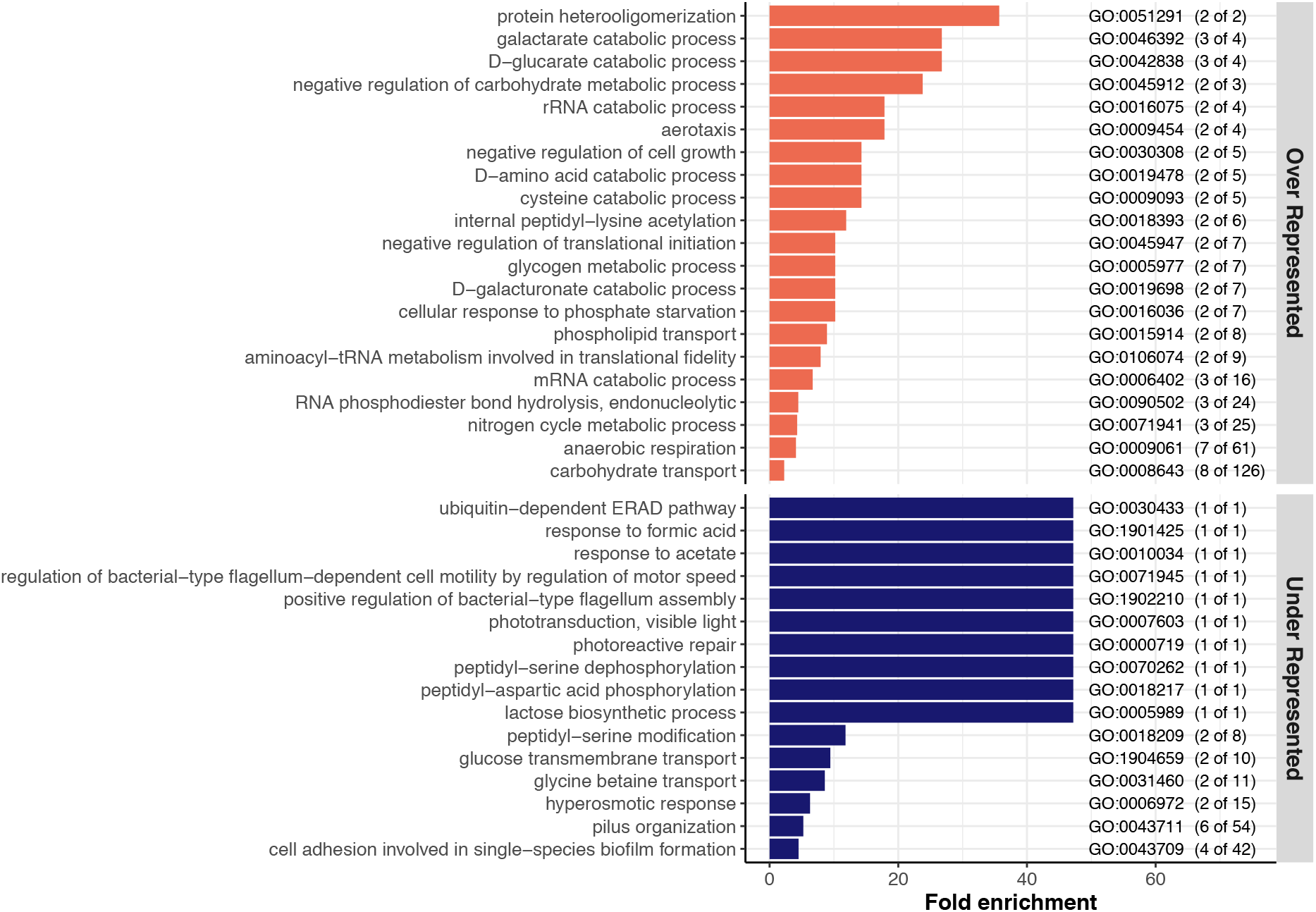
Visualization of the significantly over- and under-represented GO biological processes differentially regulated only in Ag43-mediated aggregates. Upper panel corresponds the over-represented (orange bars) pathways while bottom panel represents the under-represented pathways (blue bars). The GO enrichment analysis has been performed via PANTHER using the biological process category. The genes included in this analysis are the 243 genes commonly regulated in all the Ag43-mediated aggregates (Fig. 3 Red bar). The GO term for each biological process pathway is indicated at the right. The numbers in parentheses indicate the number of genes significantly up/downregulated in the aggregates compared to non-aggregated cells out of the total number of genes in the pathway. Note: GO:0030433 for ubiquitin-dependent ERAD pathway contains gene *ybeT* which is a prokaryotic homolog of a eukaryotic Sel1 repeat-containing protein.

**Figure 6.**
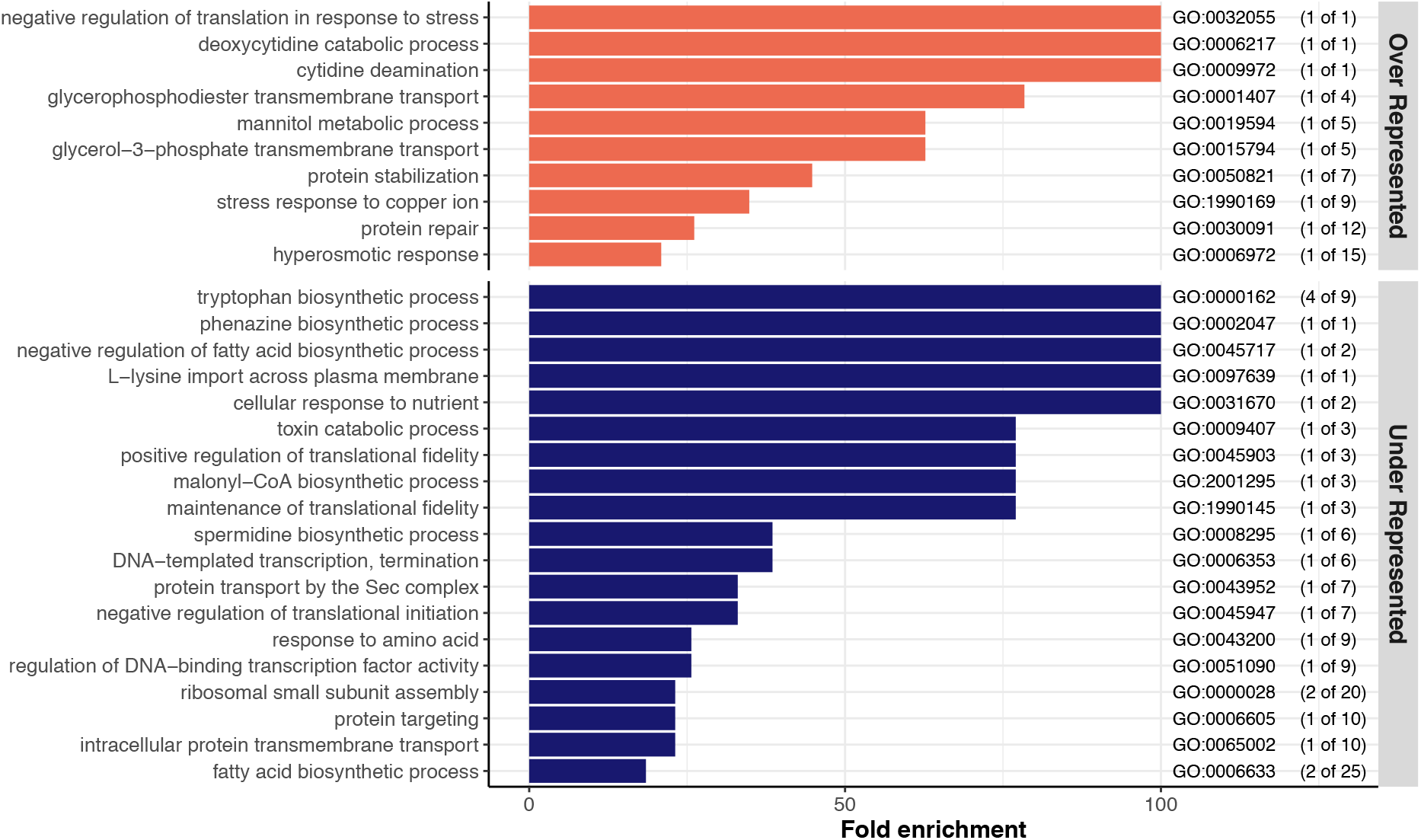
Visualization of the significantly over-versus under-represented GO biological processes differentially regulated only in Nanobodies-mediated aggregates. Upper panel corresponds to the over-represented (orange bars) pathways while bottom panel represents the under-represented pathways (blue bars). The GO enrichment has been performed via PANTHER using the biological process category. The genes included in this analysis are the 36 genes commonly regulated in all the Nanobodies -mediated aggregates (Fig. 3 Green bar). The GO term for each biological process pathway is indicated at the right. The numbers in parentheses indicate the number of genes significantly up/downregulated in the aggregates compared to non-aggregated cells out of the total number of genes in the pathway.

Beyond the specific induction of genes involved in catabolic processes of some sugar acids (*garK, garL, garR, uxaA, uxaC* for D-glucarate, galactarate, D-galacturonate) or D- and L-cysteine catabolism (*dcyD, yhaM*), genes encoding enzymes only active during anaerobiosis were also upregulated. These genes include *yhaM*, which codes for the major anaerobic cysteine-catabolizing enzyme (Loddeke et al., 2017), *ygfT* involved in the catabolic process of urate (Iwadate & Kato, 2019) and genes involved in different aspects of anaerobic respiration (*frdD, napC, nirD, glpA, torA, ynfE, dmsA*). This could indicate that Ag43-mediated aggregates display an enhanced level of anoxic conditions as compared to nanobody-mediated aggregates. Additional toxin/antitoxin genes, with notably the *mazEF* TA module, *relE* and *chpB* were also differentially expressed only in Ag43-mediated aggregates. Interestingly, the three toxins MazF, RelE and ChpB impair translation by degrading RNAs, suggesting that Ag43-mediated aggregates also display an enhanced reduction of translation. While only a few tRNA encoding genes were down-regulated in the core response to aggregation (7 genes), an important number of other tRNAs were specifically downregulated in Ag43-mediated aggregates (24 genes) (**see Supplementary Table S2 and S4**).

Finally, native aggregates are characterized by under-represented genes related to motility and surface appendages. Multiple genes related to pilus/fimbriae, known to counteract aggregation (Hasman et al., 1999; Korea et al., 2010), were downregulated (*htrE, sfmH, yadC, yadK, yehA, yehC, yehB*). The impact on motility is less clear since both *bdm*, a gene known to promote motility (Kim et al., 2015) and *ycgR*, encoding a phosphodiesterase described to impair motility (Nieto et al., 2019), were simultaneously downregulated in Ag43-mediated aggregates. By contrast, synthetic nanobody-mediated aggregation generated a limited specific response in addition to the core response. We detected the upregulation of genes encoding the small protein chaperone IbpB, the ribosome modulation factor Rmf, together with downregulation of genes encoding some ribosomal proteins (*rpsD, rpsK* and *rpsM*) that might indicate some elevated level of stress and transition to lower growth rate. We also measured a clear reduction of the genes encoding enzymes responsible for the synthesis of tryptophan (*trpA, trpB, trpC* and *trpD*).

## Discussion

The formation of bacterial aggregates has significant relevance in different types of natural environments as well as in clinical settings. To better understand the physiology of bacterial aggregates, we determined the transcriptomic response of *E. coli* to aggregation and the specificity of this response regarding the adhesin that mediates aggregation. We have shown that aggregation led to a significant and adhesin independent core transcriptome, and therefore a physiologic response in aggregated bacteria compared to non-aggregated bacteria. The transcriptional profile of the aggregated bacteria indicates a response to anoxia and other stressful conditions created by aggregation. In addition, we observed that many pathways involved in protein synthesis, DNA replication, cell division and outer membrane maintenance were strongly downregulated in aggregates, indicating of a reduced bacterial growth. Furthermore, we found that many stress resistance systems, including antibiotic tolerance and acid stress, were upregulated, correlating with the fact that aggregates can withstand stressful conditions. Finally, we also observed that many genes were differentially regulated only when the aggregates were mediated by a native *E. coli* adhesin (Ag43), essentially extending the aggregation core-response. This is indicative of a more complex and potentially time-adapted response when aggregates are formed via proteins that have been naturally present and co-evolved with their host bacterial genomes. These results are summarized in **Figure 7**.

**Figure 7.**
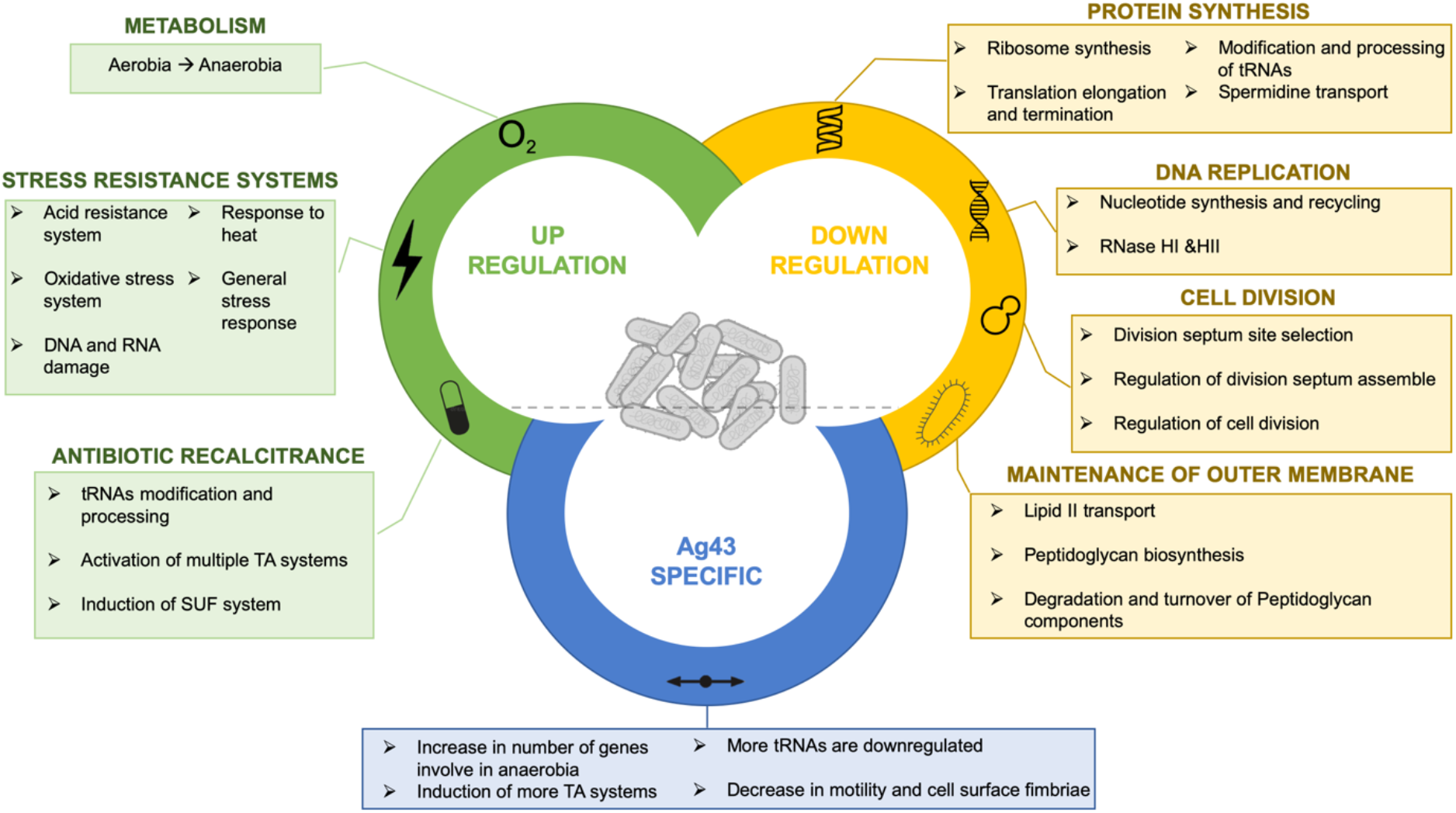
Summary of the different biological processes regulated during *E. coli* auto-aggregation. On the top are represented the main up and down regulated biological processes, in green and yellow respectively, belonging to the core response of *E. coli* towards auto-aggregation (524 genes total). At the bottom are represented the main changes specifically found in aggregates formed via the native *E. coli* adhesin Ag43 (243 genes). The genes specifically regulated within aggregates formed by synthetic adhesins are not shown in this figure.

Aggregation is often considered as an early phase of biofilm formation and aggregates themselves are fully integrated in the definition of biofilms. It has been previously shown that spontaneous 48 hour-old aggregates of *P. aeruginosa* displayed properties resembling those of biofilms, such as reduced growth rate, enhanced tolerance to antimicrobial or reduction of phagocytosis (Alhede et al., 2011). Similarly, *P. aeruginosa* 24 hour-old aggregates on alginate beads presented a transcriptional response indicating anoxic conditions (Sønderholm et al., 2017). Anoxia is also a well-known condition within biofilms formed by facultative aero-anaerobic bacteria such as *P. aeruginosa* and *E. coli* (Stewart & Franklin, 2008). This is due to the fact that the bacteria at the periphery of the biofilm actively consume oxygen, gradually reducing oxygen levels at the center of the biofilm (Stewart & Franklin, 2008). Our results indicate that anoxic conditions are also present in young (3 hour) *E. coli* aggregates. Hence, although aggregates are considerably smaller structures than biofilms, this rapid induction of anaerobia observed in our study may be due to similar mechanisms.

We and others have previously shown that one of the major physiological responses of bacteria within biofilms is the induction of stress response (Beloin et al., 2004; Schembri et al., 2003; Whiteley et al., 2001). The induction of stress response is a consequence of the drastic physico-chemical conditions prevailing with the biofilm environment. We show in this study that aggregation results in the induction of multiple stress resistance systems (e.g. acid, oxidative, heat shock, copper ion, cadmium), toxin-antitoxin systems, and starvation-related genes, along with a decrease in cellular functions related to bacteria growth. Interestingly, *P. aeruginosa* 24 hour-old aggregates on alginate beads also displayed some signs of stress and reduced growth with induction of heat-shock genes and repression of genes related to ribosomes biogenesis and functioning (Sønderholm et al., 2017). The tolerance to various stresses observed in aggregates would therefore not be directly related to the experimental conditions, timing, or species *per se*, but rather due to aggregated bacterial physiology. Although aggregates are smaller, contain fewer bacteria, and do not produce a large amount of extracellular matrix, they still possess many biofilm properties including their increased tolerance to antimicrobial agents. Interestingly, slow-growth, starvation and stresses that are encountered by aggregated cells, have for long been described as triggering aggregation (Bossier & Verstraete, 1996). This suggests that in a natural environment, these triggers might be maintained during and after formation of aggregates, therefore allowing the cells to sustain and amplify a specific physiology to resist to the surrounding stresses.

Many studies showed that bacteria are able to sense contact with abiotic surfaces and have shown that the attachment of surface structures or deformation of the cell envelope results in physiological responses (Bhomkar et al., 2010; Ellison et al., 2017; Geng et al., 2014; Persat et al., 2015; Schwan et al., 2005). These responses are mainly mediated by the activation of two-component systems and the production of secondary messengers such as c-di-GMP. While we did not detect a clear modification of the expression of two-component system sensors or regulators, we cannot exclude that some phosphorelay systems have been induced or shut down upon aggregation. Several di-guanylate cyclase or phosphodiesterase-encoding genes were down-regulated in aggregates, and while it is difficult to infer the exact net results of these regulations on c-di-GMP levels, it is possible that signaling pathways could be altered during aggregation.

Whether the transcriptional response to aggregation is due to bacterial-bacterial interaction/surface sensing or whether it is the aggregate environment that influences the transcriptome of bacteria – or both-is therefore still an open question. It is indeed possible that the contact between bacteria leads to an early response that favors aggregation by decreasing motility and the expression of other surface structures. This switch is also characteristic of early biofilm formation notably through c-di-GMP signaling (Povolotsky & Hengge, 2012). Subsequently, the environment within the aggregates would induce a secondary response including induction of anaerobic metabolism and expression of multiple stress resistance systems. Further studies will be necessary to test these hypotheses.

If bacterial-bacterial interaction signals the transcriptional shift, one can then wonder whether the type of adhesin involved in the contact influences this signaling. The enhanced transcriptional response observed when using the native adhesin Ag43 instead of the synthetic ones suggests that it is the case, beyond a core response to aggregation. This Ag43-specific response did not correspond to new functions or pathways but seemed to reinforce the pathways already up-or down-regulated in the core response to aggregation. Whether this enhanced response is a direct consequence of specific interactions mediated by Ag43 and/or a response to a different global architecture of the aggregates is unknown. Nevertheless, this suggests that the evolution that led to the generation of self-recognizing proteins was also associated to an adaptation of aggregation-mediated signal transduction. It would therefore be interesting to evaluate the transcriptomic response of aggregates mediated by other native adhesins other than Ag43, to see if we observe a similar type of response.

Like Ag43, other type of adhesins such as the Va autotransported adhesins like TibA or AIDA-I (SAATs) (Klemm et al., 2006), trimeric autotransporters (TAAs), conjugation pili (Ghigo, 2001), or fimbriae including curli or type IV pilus, can mediate aggregate formation through homotypic interactions and thus participate in kin recognition (Adams et al., 2019; Béchon et al., 2020; Clavijo et al., 2010; Collinson et al., 1996; Diodati et al., 2015; Hélaine et al., 2005; Łyskowski et al., 2011; Strassmann & Queller, 2011; Wall, 2016). Although all these cell surface appendages participate in kin recognition, they might trigger different transcriptional responses upon aggregation. Alternatively, exploring transcriptomic changes upon co-aggregation, which refers to interactions between genetically distinct bacteria (Kumar et al., 2019; Ochiai et al., 1993), that can be mediated via heterotypic interactions between SAATs or TAAs in closely related *E. coli* (Ageorges et al., 2019; Khalil et al., 2020; Klemm et al., 2006), may show different changes compared to auto-aggregation.

In conclusion, this study provides new insights into the specific properties and physiology of aggregates and opens the way to the characterization of adhesin-specific responses that could be extended to other systems for a better understanding of the functions allowing enhanced tolerance of bacteria to environmental stress.

## Material and Methods

### Bacterial strains and growth conditions

Bacterial strains used in this study are listed in **Supplementary Table S8**. All strains were grown in Miller’s Lysogeny Broth (LB) (Corning) supplemented with chloramphenicol (25μg/mL), kanamycin (50μg/mL) or ampicillin (100μg/mL) when needed. Cultures were incubated at 37°C with 180 rpm shaking during growth phase or at 37°C in static condition during aggregation phase. Cultures on solid media were done on LB with 1.5% agar supplemented with antibiotics when required. Prior to liquid cultures, bacteria were always streaked on LB agar from glycerol stocks. All media and chemicals were purchased from Sigma-Aldrich unless mentioned otherwise. All experiments and genetic construction were done in *E. coli* K12 MG1655 (F^-^,λ^-^, *rph-1)* obtained from the *E. coli* genetic stock center CGSC#6300.

### Strain construction

Insertions of the cassette carrying the chloramphenicol resistance gene and the λP_R_ or the synthetic adhesin construction were done as follows. First, *E. coli* K12 MG1655 strain was transformed with the pKOBEGA plasmid, which contains the λ-Red operon under the control of the arabinose inducible *araBAD* promoter. After inducing the expression of the λ-Red genes, electro-competent cells were prepared from this strain. In parallel, the CmPcL cassette and the synthetic adhesins construction were amplified by PCR (PCR master mix, Thermo Scientific, F548) using long floating primers that carry 40 bp of homology with the insertion region at each end. The CmPcL cassette was amplified from an existing strain in the lab collection and the synthetic adhesins were amplified from plasmid carrying the already made construction kindly sent by Ingmar H. Riedel-Kruse (Glass & Riedel-Kruse, 2018). The PCR products were then dialyzed on a 0.025 μm porosity filter and electroporated into the *E. coli* K12 MG1655 pKOBEGA strain. After electroporation, the bacteria were spread on LB plates + the appropriate antibiotic and incubated overnight at 37°C. Once the mutants were verified by PCR, the pKOBEGA plasmid was removed by plating at 42°C and the constructions were transduced using P1vir into a clean genetic background. The strains were then verified a final time by Sanger sequencing. Primers used in this study are listed in **Supplementary Table S9**.

### Aggregation curves

1 mL of overnight cultures for each strain were diluted to OD_600_ = 3 in spent LB to prevent growth during the experiment. At the beginning of the experiment, all the tubes were vortexed and 50 μL was removed (1cm below the top of the culture) and mixed with 50 μL of LB before measurement of the OD_600_. The cultures were left on the bench for 6 hours and the OD_600_ measurement 1 cm below the top of the culture was collected every hour.

### Amikacin survival assay

Bacteria were grown as described above until OD_600_ = 0.5 and then left for aggregation at 37°C for 3 hours. Bacteria were then treated with 0, 30x, 40x, 50x or 60x MIC that was previously determined (2μg/mL). The antibiotic solution was added directly into the tube on top of the bacteria and carefully mixed with the culture in order not to break the aggregates. The tubes were then put at 37°C for 18 hours. After treatment, the bacteria were washed twice and the aggregates were thoroughly vortexed in order to disrupt a maximum of aggregates. Serial dilutions from 10^−1^ to 10^−6^ were performed and spotted on LB agar, and placed overnight (14-16 hours) at 37°C before counting. Results were plotted and statistical analyses were performed using R software (version 4.0.2) implemented in RStudio (version 1.3.1093) using the *ggplot2* (Wickham, 2009) and *ggpubr* (Kassambara, 2020) packages.

### Aggregation experiment and RNA extraction

Culture sampling, RNA extraction, and RNAseq were performed in two different batches, each corresponding to the Ag43 and nanobodies experiments. Bacteria were grown as described above until OD_600_ = 0.5 and then left at 37°C for 3 hours in separating funnels in static condition (**Supplementary Fig. S2**). Then, by opening the tap of the separating funnel, 1mL of the lower part of the culture, corresponding to the aggregated part (in strains forming aggregates), was collected and placed in an Eppendorf tube. 150 μL of the aggregated or non-aggregated cells were then washed with 2 volumes of RNAprotect (Qiagen) to prevent RNA degradation, centrifugated, and the pellets were kept at -80°C until extraction. Total RNA was extracted using MP Biomedicals FastRNA Pro BLUE KIT following provider’s manual and treated with Ambion Turbo DNA-free kit to remove DNA contamination (Thermofischer Scientific; AM1907). Total RNA from 4 independent replicates for each strain/couple were quantified using the Qubit RNA HS Assay (Invitrogen) and confirmed on RNA6000 RNA chips on Bioanalyzer (Agilent) for its quality and integrity.

### Microscopy

During sampling of bacteria prior to RNA extraction, a drop of each type of aggregate was spotted on a microscopy glass (Superfrost Plus, Thermo Scientific) and covered with coverslip. The slides were then left on the bench for 15 minutes in order to let the aggregates settle. 20 pictures of each type of aggregates were then taken using epifluorescence microscope (EVOS M7000, Invitrogen). Images were then analyzed using FIJI software (Version 2.9.0). Using a macro repeating the exact same action for each image, they were smoothed to improve selection of aggregates over isolated cells, background was set to black, converted to mask and then using the analyze particle and the measure functions, the number of aggregates and their area were determined and imported into excel.

### RNA sequencing

RNA samples from the Ag43 experiment were subjected to the QIAseq Fast Select -5s/16s/23s kit (QIAGEN) for ribosomal RNA depletion according to manufacturer instructions. The NEBNext Bacteria rRNA Depletion Kit (New England Biosciences) was used for the nanobodies experiment RNA samples. No significant difference in rRNA removal efficiency was observed between the 2 sample sets. The libraries from both experiments were generated using the TruSeq Stranded Total RNA library Preparation Kit (Illumina, USA) following the manufacturer’s protocol. Library quality control was performed on an Agilent BioAnalyzer. Multiplexed RNA sequencing was performed on the Illumina NextSeq 500 platform to generate 150 bp paired-end reads. Sequencing was performed to a depth of 8 million reads per sample, for each of 16 samples per experiment (32 total).

### Bioinformatic analysis

The RNA-seq analysis was performed with Sequana (Cokelaer et al., 2017). In particular, we used the RNA-seq pipeline (v0.11.0), (https://github.com/sequana/sequana_rnaseq) built on top of Snakemake 6.1.1 (Köster & Rahmann, 2012). Briefly, reads were trimmed from adapters using Fastp 0.20.1 with default settings (Chen et al., 2018) then mapped to the Escherichia coli str. K-12 substr. MG1655 genome assembly ASM584v2, accession number GCF_000005845.2 from NCBI using Bowtie2 (version 2.4.2) (Langmead & Salzberg, 2012). FeatureCounts 2.0.1 (Liao et al., 2014) was used to produce the transcript count matrix, assigning reads to features using corresponding annotation from NCBI guided by strand-specificity information. Quality control statistics were summarized using MultiQC 1.10.1 (Ewels et al., 2016). Statistical analysis on the count matrix was performed to identify differentially regulated genes, by performing 8 different comparisons of each aggregating sample to its non-aggregating counterpart. Differential expression testing was conducted using DESeq2 1.24.0 (Love et al., 2014) indicating the significance (Benjamini-Hochberg adjusted p-values, false discovery rate FDR < 0.05) and the effect size (fold-change) for each comparison.

Raw transcript counts per gene were imported into RStudio and normalized using the vst transformation in the DESeq2 library 1.24.0 (Love et al., 2014). The principal component analysis (PCA) was calculated from vst normalized counts using prcomp() from base stats v4.0.2, and then visualized using autoplot() from the package ggfortify v0.4.13 (Tang et al., 2016). In order to compare genes commonly regulated between aggregate and non-aggregate comparisons, we extracted a gene list per comparison containing all genes where |log2FC| > 1 and padj < 0.05. These gene lists were joined and visualized using the upsetR package (version 1.4.0) (Conway et al., 2017). The significantly regulated genes common to all comparisons or unique to each experiment were extracted as gene lists and imported into the PANTHER bioinformatics web server (http://www.pantherdb.org) (Mi et al., 2021; The Gene Ontology, 2019) (**Supplementary Tables S1, S2, S4, S6**). The GO groupings that were over or under-represented in each experimental group (common to all or unique to each Ag43/nanobodies experiment) were calculated based on the number of significantly regulated genes, relative to what would be expected by chance. All processed sequencing files and R scripts used to produce the figures in the manuscript are available on Zenodo (Chekli & Stevick, 2023).

## Acknowledgements

This work was supported by grants from the French Government’s Investissement d’Avenir program, Laboratoire d’Excellence “Integrative Biology of Emerging Infectious Diseases” (grant n°ANR-10-LABX-62-IBEID) and by the Fondation pour la Recherche Médicale (grant DEQ20180339185). YC was supported by a MENESR (Ministère Français de l’Education Nationale, de l’Enseignement Supérieur et de la Recherche) fellowship. RJS was supported by a grant from the Philippe Foundation. Biomics Platform, C2RT was supported by France Génomique (ANR-10-INBS-09-09) and IBISA

## Data Availability

The raw sequencing data for this study have been deposited in the European Nucleotide Archive (ENA) at EMBL-EBI under accession number E-MTAB-11396. All other raw data, processed sequencing files, and scripts to reproduce the figures in the manuscript are available in the Zenodo repository (Chekli & Stevick, 2023).

