## Supplementary Information for "*Escherichia coli* aggregates mediated by native or synthetic adhesins exhibit both core and adhesin-specific transcriptional responses"

### Supplementary Text

We determine the Gene Ontology (GO) groupings that were over or under-represented among significantly regulated genes in the core aggregate vs non-aggregate response. Multiple pathways were found to be significantly over- (26 pathways) or under-represented (36 pathways) in aggregates compared to non-aggregate controls (**Fig. 4 and Supplementary Table S3**).

**Aggregation lead to profound metabolic changes and a global downshift of essential cellular functions** (supplementary explanation)

*Anaerobic metabolism*

Some of the over-represented pathways are linked to a switch from aerobic to anaerobic metabolism (pink highlighted pathways in **Supplementary Table S3**). For example, we found that fermentation was significantly enriched, with three genes, *hyaABC,* belonging to the type B cytochrome electron transfer, activated under anaerobia. Several anaerobic respiration chains such as the nitrate reductase A complex (NRA, *narGHIJ*)) and GlpD, that use glycerol-3-phosphate as an electron donor and nitrate as an electron acceptor (Garland et al., 1975), as well as nitrite reduction with the menaquinol-cythochrome C reductase (*nrfCD*) and the nitrite reductase complex (*nrfAB*) that reduce nitrite into ammonium, were induced upon aggregation.

*Amino acid metabolism, nucleotide production and cell growth*

In addition to some over-represented amino-acid metabolism pathways, the spermidine transmembrane transport (*potABC*) is under-represented in aggregates. Spermidine is one of the major polyamines used by *E. coli.* One of its functions is to stimulate bacterial growth, by improving protein synthesis and facilitating the normal assembly of ribosomes (Igarashi & Kashiwagi, 2006). In parallel, genes encoding for production of nucleotides, especially purines, are downregulated. In fact, four pathways involved in the synthesis and/or salvation of purines are under-represented: AMP salvage, purine nucleotide interconversion, purine nucleobase salvage, purine ribonucleoside biosynthetic process. The synthesis of pyrimidines is affected as well: dTMP biosynthetic process and pyrimidine nucleoside triphosphate metabolic process are under-represented. In addition, both RNase H genes (*rnhA* and *rnhB*), coding proteins that are involved DNA replication by preventing initiation of replication from other sites than OriC and DNA repair by removing misincorporated ribonucleotides in DNA (Vaisman & Woodgate,

2015), are downregulated in aggregates. This decrease in nucleotide synthesis genes and recycling of nucleotides as well as the downregulation of both RNase HI and HII could lead to a decrease in the fidelity and the rate of DNA replication and therefore a decrease in the growth rate of bacteria. This, therefore, indicates a trend towards some reduction of growth functions. This observation is further supported by the fact that the rewiring of several metabolic processes observed upon aggregation is also accompanied by a general slowdown of essential growth functions linked to ribosome assembly and functioning (green highlighted pathways in **Supplementary Table S3**) as well as DNA replication and division (orange highlighted pathways in **Supplementary Table S3**).

Indeed, small (*rpsU, rpsP, rpsT*) and large (*rpmA, rpmB, rpmG, rplc, rplW*) ribosomal subunit assembly pathways are under-represented in bacterial aggregates along with ribosome assembly factors and RNA helicases (*deaD* and *dbpA*). Consistently, genes involved in translation elongation and translation termination are also down regulated in aggregates, *i.e. prfA, prfB,* and *prfC* encoding for three peptide chain release factors as well as *lepA, tsf* and *yeiP* encoding three elongation factors. A decrease in release factor function can lead to an increased time to terminate/release a protein and a decreased accuracy of termination site (Baggett et al., 2017).

Three pathways involved in the regulation of cell division as well as in the positioning and assembly of the division septum are also under-represented in aggregates: division septum site selection, regulation of division septum assembly, regulation of cell division. Indeed, the *minCDE* and *slmA* genes encoding proteins involved in directing the septation to the proper site in dividing *E. coli* cell are downregulated as well as the genes *cedA* and *mioC* encoding proteins involved in the regulation of cell division (Katayama et al., 1997; Lies et al., 2015)

*Modification of cell envelope*

We observed that cell envelope synthesis and especially peptidoglycan (PG) synthesis were greatly under-represented (highlighted in violet in pathways in **Supplementary Table S3**). Among the genes included in these pathways, different enzymes involved in the transport of the building blocks that serve the synthesis of PG are downregulated: *murJ* coding for a lipid II flipase, ensuring the transport of lipid II from the cytoplasm to the periplasm*, ispU* encoding a catalase responsible of the formation of undecaprenyl pyrophosphate (UPP), that functions as the lipid carrier for bacterial cell wall carbohydrates, and *ampG* coding for a protein involved in the transport of muropeptides into the cytoplasm (Jacobs et al., 1994).

Enzymes directly involved in the synthesis of PG are also down-regulated: *mrdA* and *mrcA* code for two D,D-transpeptidases, also referred to as penicillin binding proteins (PBPs), that crosslink the peptide side chains of peptidoglycan strands to create 4->3 links. Furthermore, several genes encoding proteins involved in the degradation and turnover of PG components are also downregulated (*mltD, mltF, mepA, mepM, mepS* and *yafK)*(de Jonge et al., 1989; Dik et al., 2017; Höltje & Heidrich, 2001; Voedts et al., 2021). YafK (also called LdtF or DpaA) specifically cleaves the cross-link of Lpp, the Braun's lipoprotein, to the PG. Lpp is essential to maintain membrane integrity by linking the outer membrane with PG (Leduc et al., 1992; Suzuki et al., 1978). Interestingly, the *lpp* gene itself was upregulated in aggregates suggesting that attachment of the outer membrane to the PG could be strengthened upon aggregation.

*c-di-GMP regulation*

Interestingly, some genes involved in c-di-GMP regulation are also modulated in aggregated cells. *yeaP* (*dgcP*) and *yliF* (*dgcI*) encoding di-guanylate cyclases, *yliE* (*pdeI*) encoding a dual di-guanylate cyclase/phospho-diesterase, and *yahA* (*pdeL*) encoding a phosho-diesterase are all downregulated. The net results of c-di-GMP regulation are difficult to infer since YahA (PdeL) has been shown to inhibit motility (Yilmaz et al., 2020) while YliE (PdeI) activity rather increases motility (Reinders et al., 2016). Meanwhile, YeaP has been linked to curli expression at transcriptional or post-transcriptional levels, but its effects are barely detectable since it is lowly expressed (Sommerfeldt et al., 2009). Some regulation of c-di-GMP related enzymes is thus probably at play during aggregation. However, the complexity of the regulation of their target genes prevents definitive conclusion on the impact of their modulation.

**Aggregates are subjected to and transcriptionally respond to different stresses**

Many genes related to various stress responses are induced upon aggregation (pathways highlighted in red in **Supplementary Table S3**). Two out the four acid resistance systems, GadABC and CadAB/LcdC, are enriched in aggregates. Each system has two components: a cytoplasmic amino acid decarboxylase and an inner membrane substrate/product antiporter (Foster, 2004). Both transport reactions produce $CO_2$ and consume a proton, which raises intracellular pH and helps *E. coli* to fight acidic stresses (Kanjee & Houry, 2013). Moreover, we found that *yhiM,* encoding YhiM, a protein necessary for glutamine and lysine-dependent acid resistance (Nguyen & Sparks-Thissen, 2012), was also upregulated in aggregates compared to non-aggregated cells, along with other acid stress response genes (*hdeAB, hchA, dps*). At low pH HdeA and HdeB, two periplasmic chaperones, bind to acid-denatured proteins (Gajiwala & Burley, 2000; Kern et al., 2007). Upon pH increase, HdeA and HdeB release bound proteins under their folded conformation (Tapley et al., 2010). The chaperone Hsp31 (HchA), plays the same role in the cytoplasm (Quigley et al., 2004).

Several genes associated to oxidative stress like hydrogen peroxide (*ychH,ygiW* and *katG)* and cellular detoxification (*katG, sodB, hcp* and *frmA)* or protection (*dps*) are significantly over-expressed in aggregates. KatG is involved in the degradation of superoxide radicals, which are byproducts of aerobic metabolism, together with superoxide dismutase (SOD) SodB. *E. coli* possesses three different SOD with complex regulation. Up-regulation of *sodB,* which is the only SOD present under anaerobic conditions (Hassan & Fridovich, 1977; Kargalioglu & Imlay, 1994), further suggests that some bacteria within the aggregates respond to an aerobic environment. In addition, we found that some other genes involved in oxidative stress were up-regulated in aggregates such as *clpA, clpB, acnA and wrbA,* which strengthens the hypothesis that certain bacteria within the aggregates maintain an aerobic metabolism.

Beside acidic or oxidative stress responses, we found that the methylglyoxal degradation III pathway (*dkgA, yqhD, gldA and hchA)* and RNA repair pathways (*hchA and rtcB)* were enriched in aggregates. Methylglyoxal is highly toxic and stressful for bacteria most likely because of its ability to interact with both protein side chains and nucleophilic centers of macromolecules such as DNA (Ferguson et al., 1998; Kalapos, 1999; Richarme et al., 2017).

### Supplementary Figures

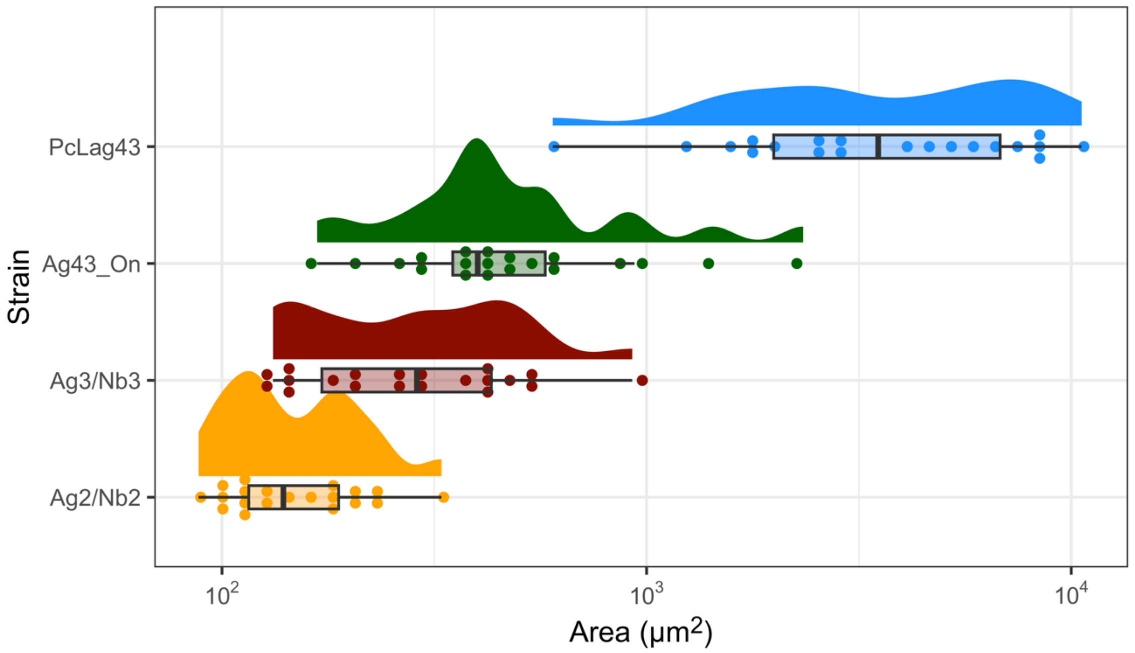


**Supplementary Figure S1. Size of bacterial aggregates.** Raincloud plot representing the average area of the aggregates formed through the native and synthetic adhesins. For each strain, 20 fields were captured, each containing multiple aggregates. The mean area of the aggregates per field is plotted in μm$^2$.

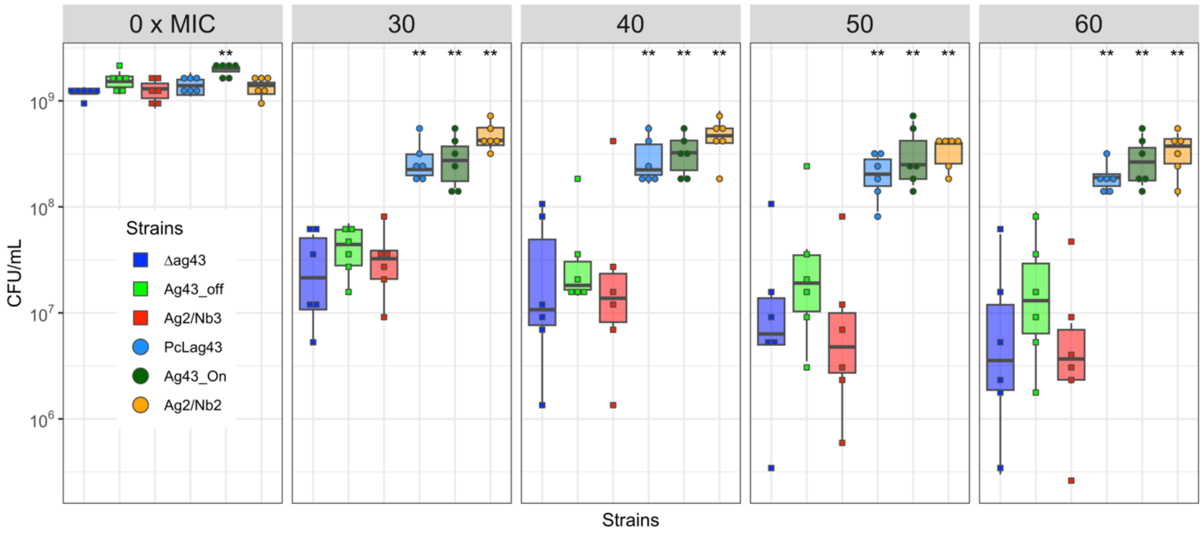


**Supplementary Figure S2. Ag43- and nanobodies-mediated aggregation leads to enhanced survival when treated with lethal concentration of amikacin.** Boxplot representing the CFU/mL for each strain after treatment for 18h with 0, 30x, 40x, 50x and 60x (each panel) the MIC of amikacin (2µg/mL). For each strain 6 biological replicates (each replicate is the mean of 2 technical replicates) have been performed, represented as a point on the graph. Statistical analysis was performed using the non-parametric Wilcoxon test for each strain compared with the Δ*ag43* non-aggregative control (**: p-value <= 0.01).

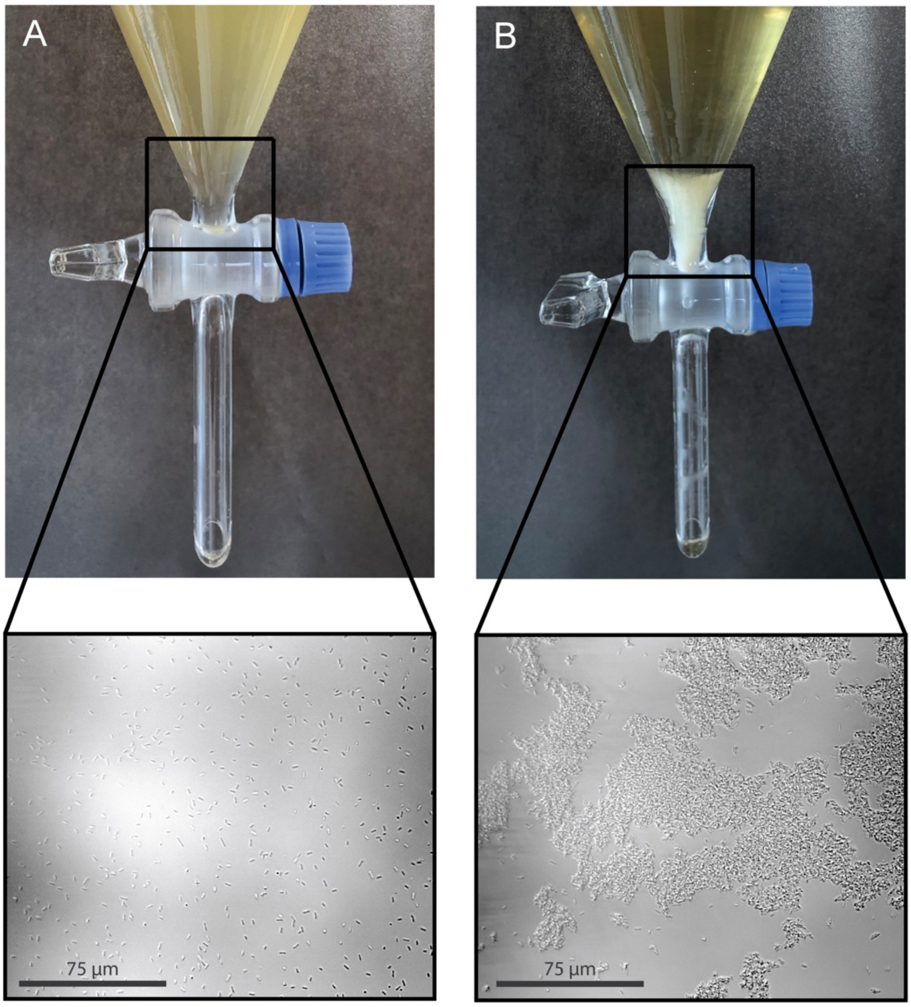


**Supplementary Figure S3. Bacterial aggregates are recovered through a separating funnel system and analyzed with optical microscopy.** (A) Example of a separating funnel containing a 3h culture of the Ag43_Off culture showing no aggregates. (B) Example of a separating funnel containing a 3h culture of the Ag43_On culture showing aggregated bacteria. Scale bar = 75 µm.

### Supplementary Tables

**Supplementary table S1. Pan response to aggregation**

**Supplementary table S2. Core response to aggregation**

**Supplementary table S3. Enrichment analysis by PantherD of the over- and under-represented pathways from the core response to aggregation**

**Supplementary table S4. The specific response to aggregation mediated by the native adhesin Ag43**

**Supplementary table S5. Enrichment analysis by PantherD of the over- and under-represented pathways from the response to Ag43-specific aggregation**

**Supplementary table S6. The specific response to aggregation mediated by the synthetic adhesins (nanobodies)**

**Supplementary table S7. Enrichment analysis by PantherD of the over- and under-represented pathways from the response to nanobody-specific aggregation**

**Supplementary table S8. Strains and plasmid used in this study**.

| Strain | Description | Source |
|---|---|---|
| *E. coli* MG1655 | $F^-$,$\lambda^-$, *rph-1* | *E. coli* genetic stock center CGSC#6300. |
| MG1655 Δ*flu* | MG1655 deleted of *agn43*, KmR. | (Chauhan et al., 2013) |
| MG1655 *PcLflu* | $\lambda P_r$ placed in front of *agn43*, resulting in constitutive expression of the gene, CmR. | This study |
| MG1655 *EndfluLacZzeo* | *lacZ* gene is transcriptionally fused at the end of *agn43*, allowing the tracking of the ON/OFF status of the natural promoter of *agn43,* ZeoR. | (Chauhan et al., 2013) |

| | | |
|---|---|---|
| MG1655 Δ*flu PcLAg2* | The synthetic adhesin Ag2 is placed at the *agn43* locus site, under the control of the $\lambda P_r$, KmR, CmR. | This study |
| MG1655 Δ*flu PcLNb2* | The synthetic adhesin Nb2 is placed at the *agn43* locus site, under the control of the $\lambda P_r$, KmR, CmR. | |
| MG1655 Δ*flu PcLAg3* | The synthetic adhesin Ag3 is placed at the *agn43* locus site, under the control of the $\lambda P_r$, KmR, CmR. | |
| MG1655 Δ*flu PcLNb3* | The synthetic adhesin Nb3 is placed at the *agn43* locus site, under the control of the $\lambda P_r$, KmR, CmR. | |
| pKOBEGA | pSC101 thermosensitive (replicates at 30°C), *araC*, arabinose-inducible λredγβα operon, AmpR | (Chaveroche et al., 2000) |
| pDSG419 | pSB3K3_TetR_pTet_Neae2v1_A4_EPEA. Plasmid carrying the Ag2 construct, KmR GenBank: MH492413 | (Glass & Riedel-Kruse, 2018) |
| pDSG375 | pSB3K3_TetR_pTet_Neae2v1_N4-1_antiEPEA-EPEA1. Plasmid carrying the Nb2 construct, KmR. GenBank: MH492440 | |
| pDSG360 | pSB3K3_TetR_pTet_Neae2v1_A8_P53TA. Plasmid carrying the Ag3 construct, KmR. GenBank: MH492448 | |
| pDSG321 | pSB3K3_TetR_pTet_Neae2v1_antiP53TA. Plasmid carrying the Nb3 construct, KmR. GenBank: MH492393 | |

**Supplementary table S9. Primers used in this study.**

| **Name** | **Sequence 5’-> 3’** |
|---|---|
| CmPcLFlu_L5 | accggcttttttattcaccctcaattcgctcaagttagtaattctcaccaataaaaaacg |
| CmPcLFlu_L3 | agcaggtattcagatgtcgtttcatcagcttttccttagatagtacatgcaaccattatc |
| L3_CmPcLAg_NB | cggtcatgatgaccgggaccacagagaggcgatggttctggtgtctcaaaatctctgatg |
| L5_CmPcLAg_Nb | gcggtgataatggttgcatgtactatctaaggaaaagctgatgattactcatggttgtta |
